# The AvianMetaNetwork: biotic interactions among birds of the continental United States and Canada

**DOI:** 10.64898/2026.05.11.723238

**Authors:** Phoebe Lehmann Zarnetske, Patrick Bills, Kelly Kapsar, Lucas Mansfield, Emily G. Parker, Caroline Roche, India Hirschowitz, Giovanni DePasquale, Sara Zonneveld

**Affiliations:** Department of Integrative Biology, Michigan State University, USA; Ecology, Evolution, and Behavior (EEB) Program, Michigan State University, USA; Institute for Biodiversity, Ecology, Evolution, and Macrosystems (IBEEM), Michigan State University, USA; Institute for Cyber-Enabled Research, Michigan State University, USA; Independent Researcher (Ornithology), Exeter, United Kingdom

## Abstract

All organisms interact with other organisms, directly, and indirectly through different ecological relationships involving multiple types of interactions. Yet at broad continental scales, we lack comprehensive information on biotic interactions, which has hindered our ability to answer macroecological and eco-evolutionary questions across scales and to fully quantify the diversity of biotic interactions as an important dimension of biodiversity. Here, we help fill these gaps with an open and comprehensive dataset and data workflow of 25,907 pairwise, directional interspecific interactions among birds spanning a continental scale. All data are empirically documented and comprise bird-bird interactions across both breeding and non-breeding ranges of 731 focal avian taxa, covering all birds in the focal region of Canada and the continental United States, including Alaska. These data also include 1,258 additional avian taxa interacting with the focal taxa outside the focal region, resulting in 1,989 avian taxa altogether. The continental scale and breadth of interspecific interactions within these data fill fundamental knowledge gaps and enable scientists and practitioners to address a myriad of questions at broader scales than were previously possible.

## Background & Summary

Biotic interactions are essential linkages among organisms that ultimately scale up to affect the functioning of entire ecosystems ^1–6^. Comprehensive knowledge of biotic interactions among organisms can greatly improve understanding and prediction of the structure and function of ecological systems, especially from local to continental scales and across time ^7–9^. While numerous studies have documented how organisms interact, studies are often local in extent, collected for specific field or experimental projects, or focused on a relatively small subset of interaction types or taxa ^10,11^, limiting their potential to provide robust data for understanding the causes and consequences of ecological patterns from organisms to ecosystems across scales ^12–14^. Analysis-ready biotic interaction data for entire classes of organisms at continental scales can help fill important gaps in knowledge to advance both fundamental and applied research.

Data on biotic interactions are critical for understanding and predicting the processes underlying species distributions, community assembly, eco-evolutionary dynamics, invasive species’ success, biodiversity patterns, and ecosystem structure and function ^5,15–23^. In addition, biotic interaction data help fill the Eltonian Shortfall knowledge gap in how species interact ^10,11,13^, and contribute to greater understanding of the diversity of biotic interactions as a unique facet of biodiversity ^22,23^. Whereas numerous macroecological studies have focused on taxonomic, trait-based functional, and phylogenetic diversity, the diversity of biotic interactions has received far less attention due to lack of sufficient interaction data at broad spatial scales ^23,24^.

Given that biotic interactions are known drivers of species distribution, success and ecosystem structure, a thorough understanding of interactions is also critical in informing species’ threat status and approaches to conservation, management and restoration goals, including achieving 30 × 30 Global Biodiversity Framework targets ^6,7,25,26^. Yet we lack the data to understand and predict how biotic interactions and the networks they form respond to past, present, and future global changes across large geographic regions and broad taxonomic groups ^12,13,23,27,28^. While there is a large amount of literature and natural history knowledge on how species interact, the interaction data that exist are often from local observational or experimental studies of a relatively small subset of interacting species ^10,12,13^, only cover a single type of interaction (typically trophic) ^29–31^, or rely heavily on inferred (non-empirical) interactions ^32^. This lack of coverage of empirically-based biotic interactions hinders our ability to evaluate how local biotic interactions scale up to affect patterns of biodiversity at broader spatial and temporal scales.

We also lack open, efficient, and reproducible workflows to synthesize the vast literature necessary to create comprehensive, empirically-based, validated, and open analysis-ready data on biotic interactions across taxa, hindering many research investigations, especially at broader scales ^12,33^. Data literacy and best practices in data science and workflow design are key to successfully addressing questions across scales ^14^. The sheer effort necessary to compile observed interactions across wide geographic areas and taxa, and from disparate sources, has hindered progress and is a main reason why biotic interactions are left out of species distribution models and analyses of environmental change impacts ^34,35^. As a result, many models of species distributions and responses to global changes leave out or implicitly assume the intricacies of biotic interactions, leading to incomplete and potentially less robust predictions at any given scale, let alone across scales ^19,36–39^. With scalable, open, and reproducible workflows constructed according to the FAIR data principles ^33,40,41^, it is possible to compile comprehensive biotic interaction data, generate metanetworks of potential interactions across many different taxa, and improve model performance ^42^.

In addition to the fields of biodiversity science and conservation biology, the growing fields of continental scale biology and macrosystems biology would especially benefit from comprehensive biotic interaction data ^14^. Interaction data involving numerous taxa that occur across wide geographic areas can provide fundamental digital information for scientists and practitioners to address numerous basic and applied questions across varying spatial and temporal scales, especially when combined with data on species spatiotemporal distribution and environmental variables ^14^. For example: understanding and predicting scaling relationships and emergent properties of ecological networks over space and time, clarifying the generality of integrative animal behavioral research across species involved in certain interaction types, and explaining how and why individuals and communities behave in certain ways given the broader context of their numerous species interactions.

A key to addressing these knowledge gaps in environmental and ecological research is providing empirically based, comprehensive, and readily usable digital natural history data on interactions among taxa at a continental scale. Birds are ideal taxa to help fill this gap; they participate in numerous positive, negative, and neutral interactions ^43–45^ which contribute to important ecosystem services and functions on every continent ^46^. Here we introduce AvianMetaNetwork, a dataset of interspecific bird interactions at a continental scale, along with an accompanying workflow and vignette to generate, summarize, and explore the data.

AvianMetaNetwork contains 25,907 documented pairwise, directional interactions among 731 focal taxa occurring in the conterminous United States, Alaska, and Canada. With 1,258 additional taxa interacting with the focal taxa, these data capture interactions involving 1,989 avian taxa, including 14.59% of the world’s 11,145 bird species ^47^. These data include 6 main types of interactions (trophic, commensalism, parasitism, competition, mobbing, facilitation; Figure 1, Table 1), and 23 specific interaction sub-types (including 3 neutral or unknown direction relationships like co-occurrence). Interaction sub-types between pairs of species include different forms of competition (e.g., over food, space), parasitism (e.g., brood parasitism, kleptoparasitism), facilitation (e.g., mixed flocking), commensalism (e.g., cavity nest creation), and mobbing (i.e., predator deterrence). There are 18,419 unique pairwise interactions in these data, since the data contain multiple documented cases of the same pairwise species interactions (Figure 1) that span 123 families in 32 orders (Figure 2).

**Table 1.** Interaction types, values, and sub-types. The interaction type contains one or more sub-types of interactions which have specific interaction effects on species.

| Interaction Type | Interaction Effects | Interaction Sub-Type | Interaction Definition |
| --- | --- | --- | --- |
| Trophic | -1,1 | Predation | Predator eats prey; predator = 1, prey = -1. |
| Competition | -1,-1 | Competition | Competing over space, territory, food, etc.; both species = -1. |
| Facilitation | 1,1 | Facilitation | When both interacting species benefit (if there is no specific facilitation interaction), e.g., creching, mutual allopreening; both species = 1. |
| Facilitation | 1,1 | Facilitation-mixed flocking | Special case of facilitation when both species benefit by being in a group; both species = 1 |
| Facilitation | 1,1 | Facilitation-comigration | Special case of facilitation when species migrate together; both species = 1 |
| Facilitation | 1,1 | Facilitation-feeding/foraging | Special case of facilitation when both species feed together, either in mixed-flocks or feeding aggregations; both species = 1 |
| Facilitation | 1,1 | Communal nesting | Special case of facilitation when species nest together, usually during breeding season; both species = 1. |
| Facilitation | 1,1 | Communal roosting | Special case of facilitation when species roost together, typically during non-breeding season; both species = 1. |
| Commensalism | 0,1 | Commensalism | One interacting species benefits and the other receives no effect (example: cavities are constructed by primary cavity nesting bird like a woodpecker, and later used by another bird for its own nest (without any conflict)); species that benefits from the interaction = 1; species that provides the benefit = 0. |
| Commensalism | 0,1 | Commensalism-scavenge | Special case of commensalism when a species consumes/scavenges a previously deceased species or deserted/addled eggs; scavenger=1, dead species=0. |
| Commensalism | 0,1 | Commensalism-chick adoption | Special case of commensalism when one species adopts another species' chicks/eggs; species that is adopting = 0, species that is adopted = 1. |
| Commensalism | 0,1 | Commensalism-call mimicry | Special case of commensalism when one species copies the call/song of another; species that is mimicking = 1, species that is being mimicked = 0. |
| Parasitism | 1,-1 | Kleptoparasitism | Special case of parasitism when one species steals food or nest material from another species; species that steals and benefits = 1, species that is stolen from = -1. |
| Parasitism | 1,-1 | Brood parasitism | Special case of parasitism when one species places its egg in another species' nest so the nesting species has to raise a conspecific (nest parasitism - includes egg-dumping); species that lays egg in another species nest = 1, species that has to care for the conspecific young = -1. |
| Parasitism | 1,-1 | Nest takeover | Special case of parasitism when one species takes over (steals) the nest of another species, sometimes kicking out the existing eggs; species that takes over = 1; species whose nest is stolen = -1. |
| Amensalism | 0,-1 | Amensalism | One species causes harm to another species without any cost or benefits to itself (e.g. flushing from nest in response to perceived predator, accidental destruction of eggs); harmful species = 0, harmed species = -1. |
| Mobbing | 0,-1 | Mobbing | Usually indicative of predation, where the smaller species mobs the bigger species; the species doing the mobbing (usually potential prey) = 0; the species receiving the mobbing = -1 (the potential predator). The interaction is coded this way because mobbing is almost always deterring a predator, which reduces the likelihood of predation. |
| Other | 0,0 | Co-occurrence | Not an interaction per se. Species occur together but it is not clear what the interaction is; each species = 0. |
| Other | 0,0 | Play | Species play with each other; both species = 0. |
| Other | 0,0 | Hybridization | When species successfully mate and produce hybrid offspring; includes intergrades (hybrids of subspecies); both species = 0. |
| Other | 0,0 | Copulation | Unsuccessful mating attempt between two individuals of different species; both species = 0. |
| Other | 0,0 | Courtship | One species attempts to court another; both species = 0 |
| Other | 0,-1 | Accidental Killing | when one species accidentally kills another, but has no intent on doing so (e.g., no intent on preying on the dead species or competing with the dead species); species killed = -1, species who causes the death = 0 |

**Figure 1.**
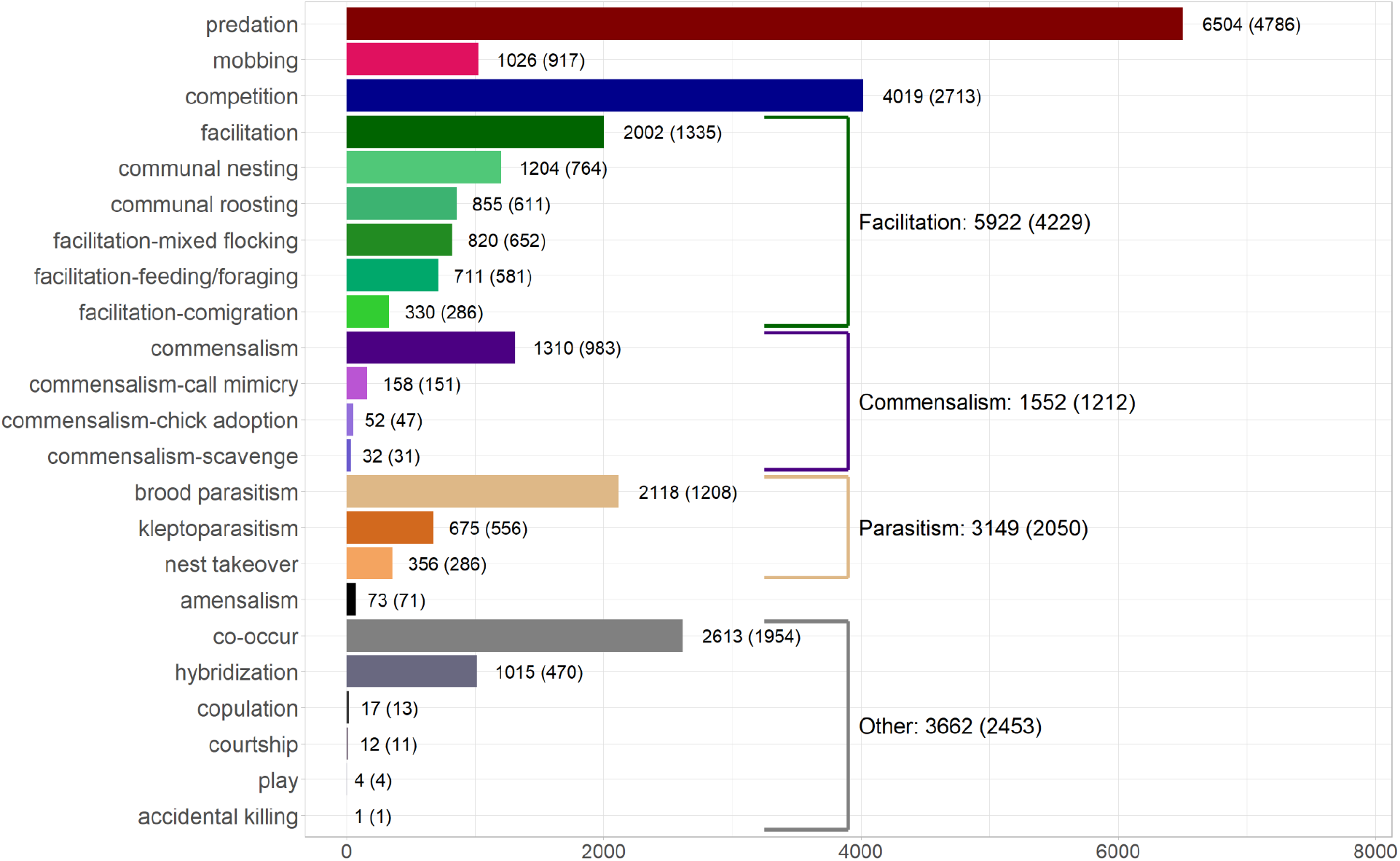
Distribution of total pairwise interactions in AvianMetaNetwork by interaction sub-type, grouped by interaction type. The first number includes instances where the same pairwise interaction is documented in multiple sources; the number of unique pairwise interactions per interaction type are indicated in parentheses (see “8_summary_vignette” described in *Usage Notes* for obtaining the total number of interaction records by interaction type and sub-type). “Unique pairwise interactions” refers to the unique combination of interaction type or sub-type and interaction effect for a given pair of taxa. Interaction type color scheme is as follows: Red: Trophic (Predation); Pink: Mobbing; Dark Blue: Competition; Greens: Facilitation; Purples: Commensalism; Oranges: Parasitism; Black: Amensalism; Grays: Other.

**Figure 2.**
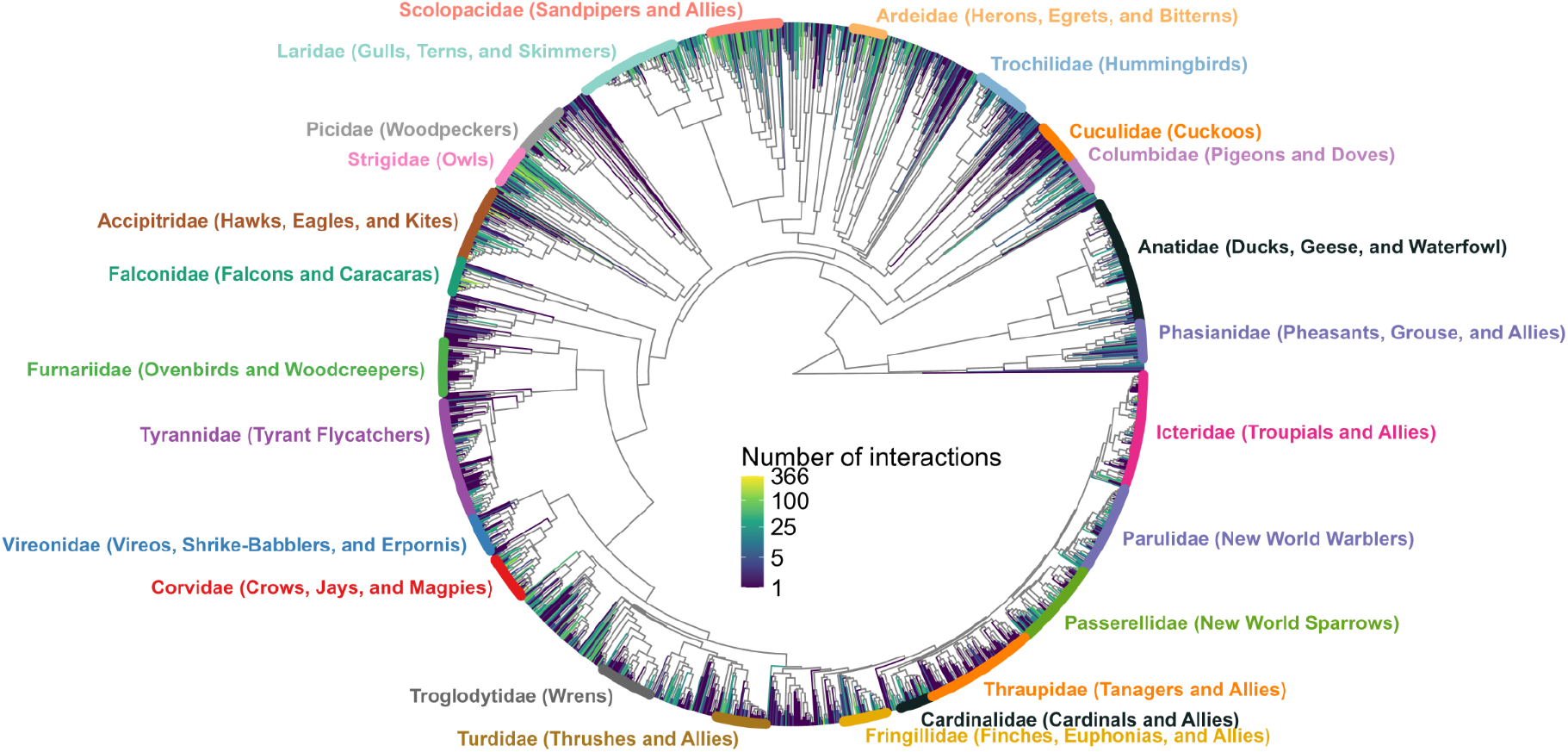
Avian phylogenetic distribution of the number of unique pairwise interactions at the species-level in AvianMetaNetwork. Branch color represents the number of interactions, with darker-branched species participating in more documented interactions. Colored tip labels represent families in the database with 25 or more total species. Phylogeny from ^54^, visualized with ^55,56^.

Advances made possible by these data and workflow include: (1) helping to fill the Eltonian shortfall knowledge gap in species interactions ^10,11,13^, (2) the ability to uncover the structure and function of species interaction networks across space and time and test multiple ecological theories involving species interactions at scales previously not possible ^14,48–52^, (3) informing conservation and management decisions across scales of biological organization, from species to ecosystems, including quantifying the influence of keystone species in their networks ^8,45,53^, and (4) providing a robust, open, and generalizable workflow for generating analysis-ready directional relationship data. With ongoing efforts to expand the database across the globe, these data will enable numerous basic and applied research and conservation investigations involving some of the most ubiquitous and extraordinary taxa across Earth’s ecosystems.

## Methods

### Methods Overview

AvianMetaNetwork contains a comprehensive compilation of documented bird-bird interactions across the continental United States (including Alaska) and Canada. Trained data recorders extracted interactions manually by systematically reviewing avian species accounts in Birds of the World ^57^, an expert curated set of species accounts for all birds ^57^. The reproducible data cleaning workflow and collaborative coding with GitHub ensure that all changes to the data (e.g., taxonomy changes, interaction interpretations) are traceable and documented. To organize levels of data cleaning, we followed the Environmental Data Initiative’s (EDI) thematic standardization for Level 0 (raw), 1 (cleaned), and 2 (analyzed) data ^58^. To encourage open and accessible data ^59^, the database and its accompanying workflow align with the Findable, Accessible, Interoperable, Reusable (FAIR) data principles for open science ^40^. The workflow used to generate AvianMetaNetwork is described in detail below and summarized in Figure 3.

**Figure 3.**
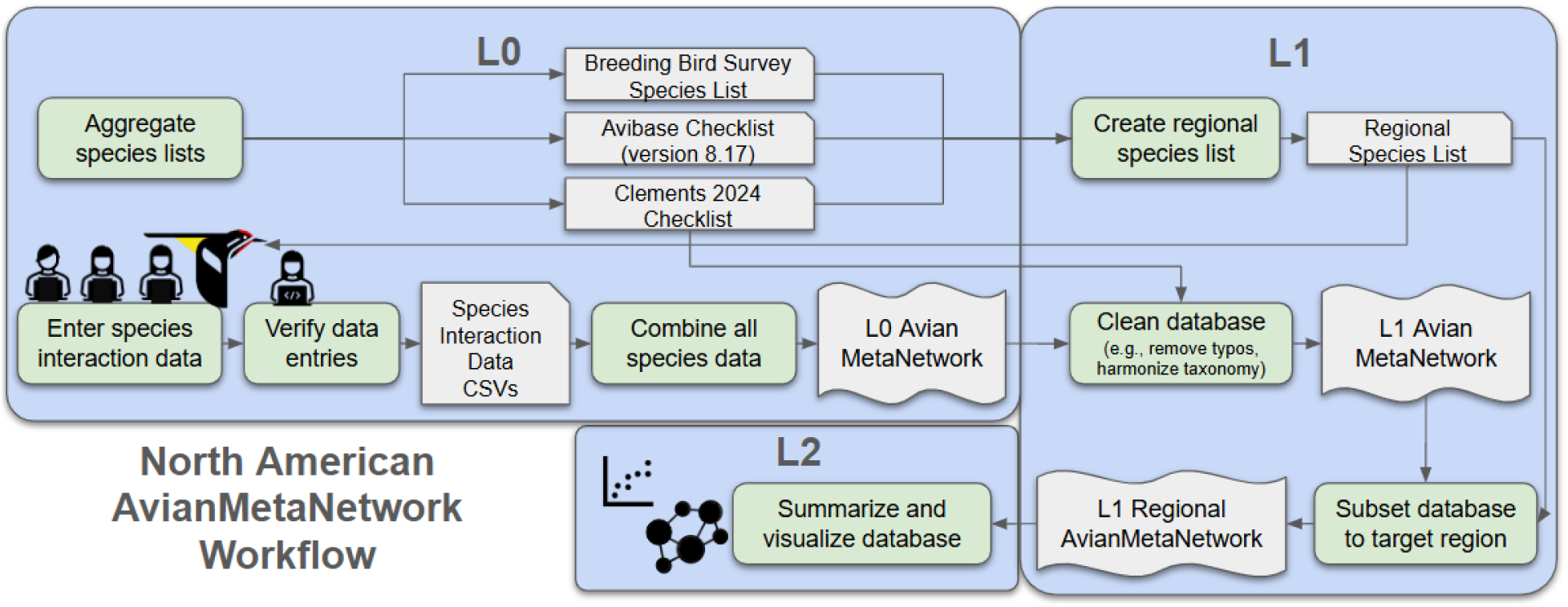
Workflow diagram for the creation of the AvianMetaNetwork. Steps are shown in green boxes. Gray boxes indicate data sets (i.e., species lists, interaction data); L0 = level 0 (raw) data; L1 = level 1 (cleaned data); L2 = level 2 (summarized data in analyses).

**Figure 3.**
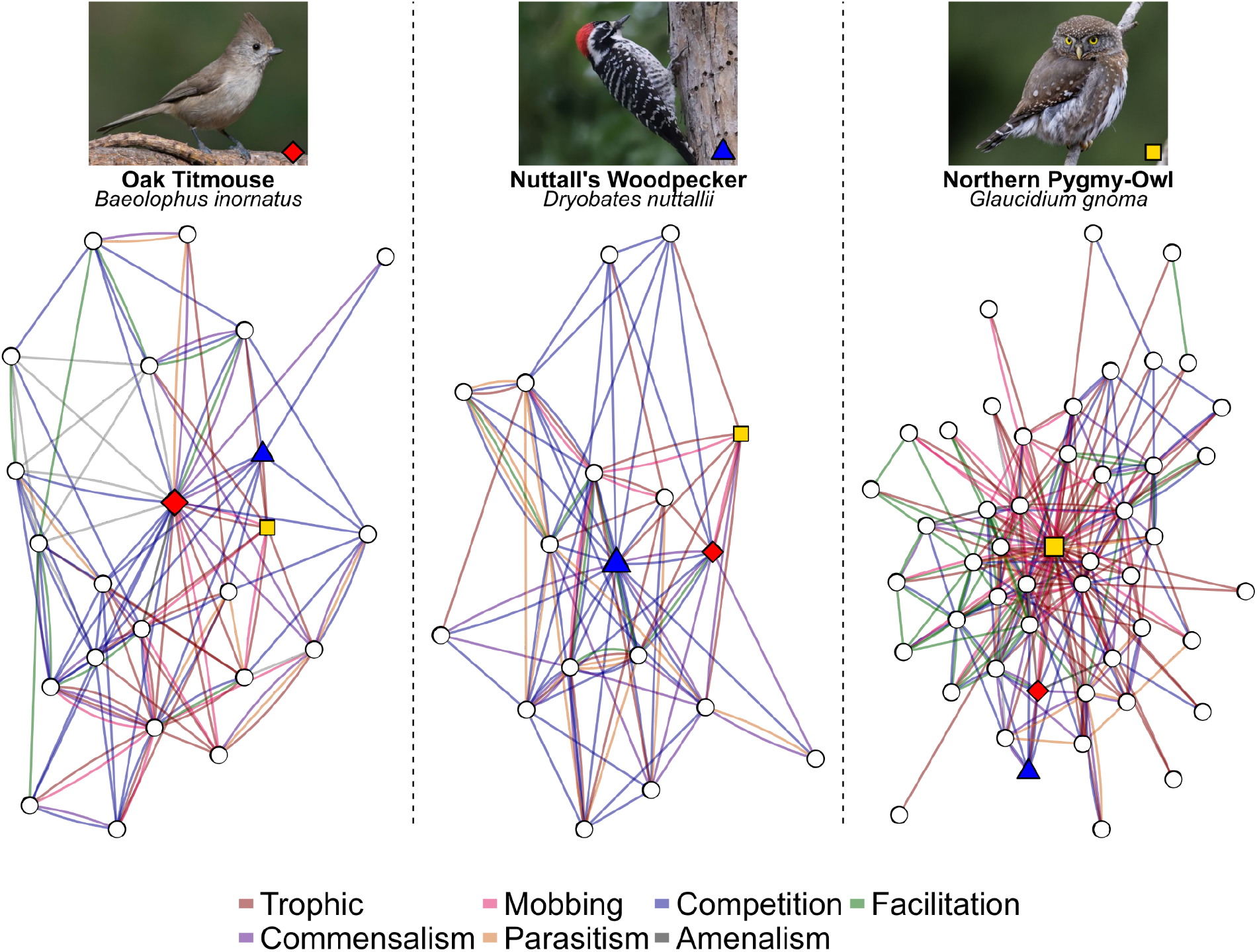
Example networks generated from AvianMetaNetwork, for the three focal species: Oak Titmouse (*Baeolophus inornatus*, red diamond), Nuttall’s Woodpecker (*Dryobates nuttallii*, blue triangle), and Northern Pygmy-Owl (*Glaucidium gnoma*, yellow square). The networks represent the focal species and all of their interacting species. Each focal species is represented on all three networks with its corresponding symbol, and other interacting species are represented with white circles. Line color designates interaction type. See *Data Code and Availability* for network plotting code. Pictures obtained from Avicommons (Oak Titmouse: Adam Jackson | CC0 2.0; Nuttall’s Woodpecker: guyincognito | CC BY-NC 2.0; Northern Pygmy-Owl: Liam Hutcheson | CC BY-NC 2.0).

### Taxa Lists

To generate our “Regional Species List” of focal species found in Canada and the continental United States for systematic review, we merged Avibase v8.17 regional checklists for “Canada”, “US lower 48”, and “Alaska” ^60–62^ with the North American Breeding Bird Survey (BBS) 2024 species list ^63^. We used the Clements/eBird 2024 checklist as the taxonomic authority for naming conventions for all taxa. To facilitate reuse of the network data, we include a “taxa1_clements” and “taxa2_clements” column in AvianMetaNetwork. This column contains the finest taxonomic resolution information available for the interaction. This column can be matched to the relevant Clements/eBird 2024 checklist scientific name in a master checklist of all taxa in AvianMetaNetwork (including both focal and non-focal interacting taxa) which is part of the Environmental Data Initiative data package (see *Taxonomic Harmonization* below for more detail). In total, there are 731 focal taxa in AvianMetaNetwork (including subspecies, genus-level taxa, hybrids and uncertain taxa) which occur in the continental United States (including Alaska) or Canada. Through the data entry process, recorders discovered 1,258 additional avian taxa that interact with focal taxa; see *Data Entry* below for more detail).

### Data Entry

To enter interactions for a given focal species, recorders systematically read through a species’ account in BOW to interpret information (Table 2). Whenever another avian taxa was mentioned in the BOW main text or figure captions, either as a common name or scientific name, recorders determined whether the text met the following criteria for a bird-bird interaction description: at least two different avian taxa are involved in an interaction (as defined by Table 2). Occasionally interactions with a focal species involved avian taxa at a finer or coarser taxonomy than species-level (e.g., subspecies or genus).

**Table 2.**
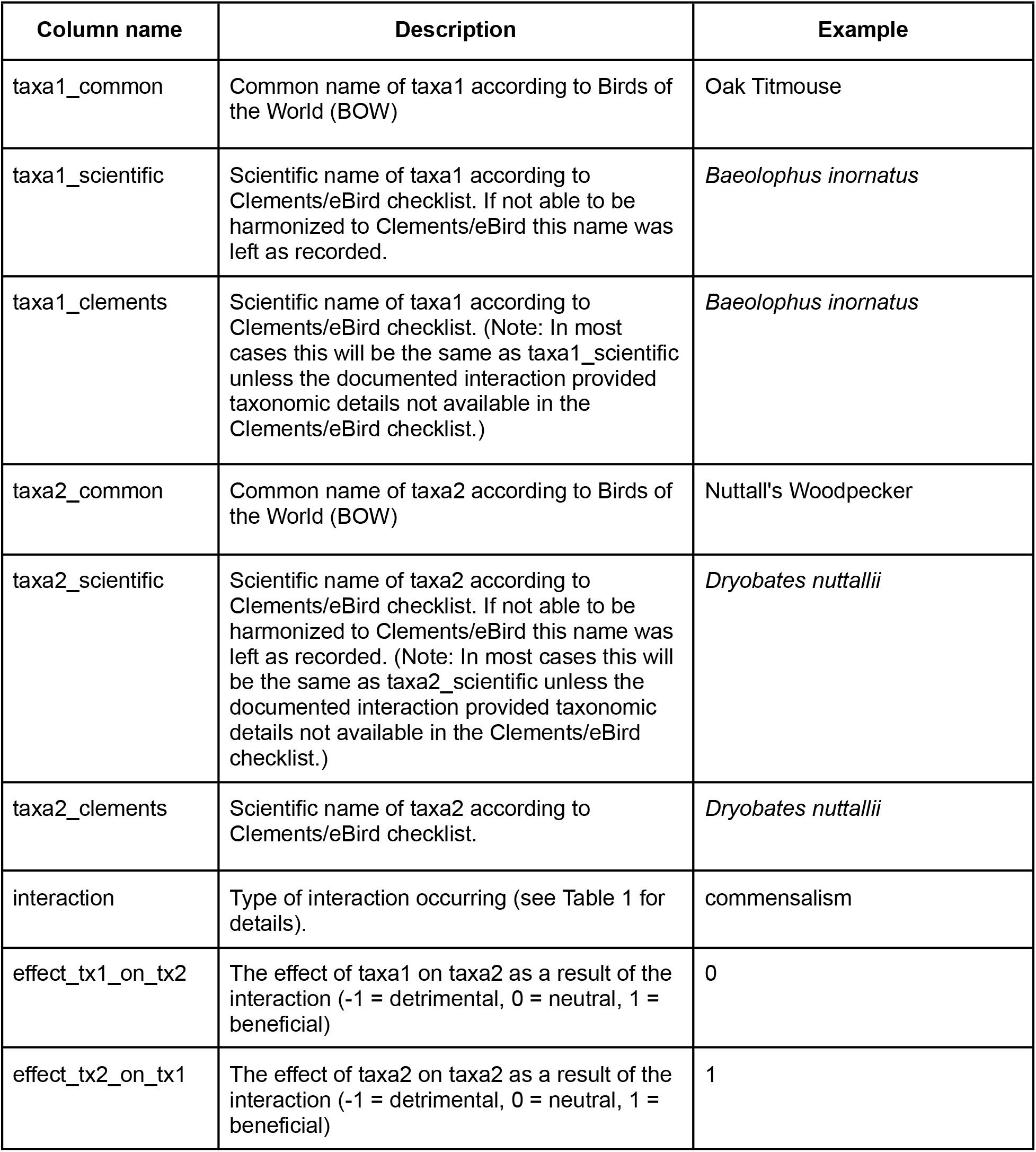
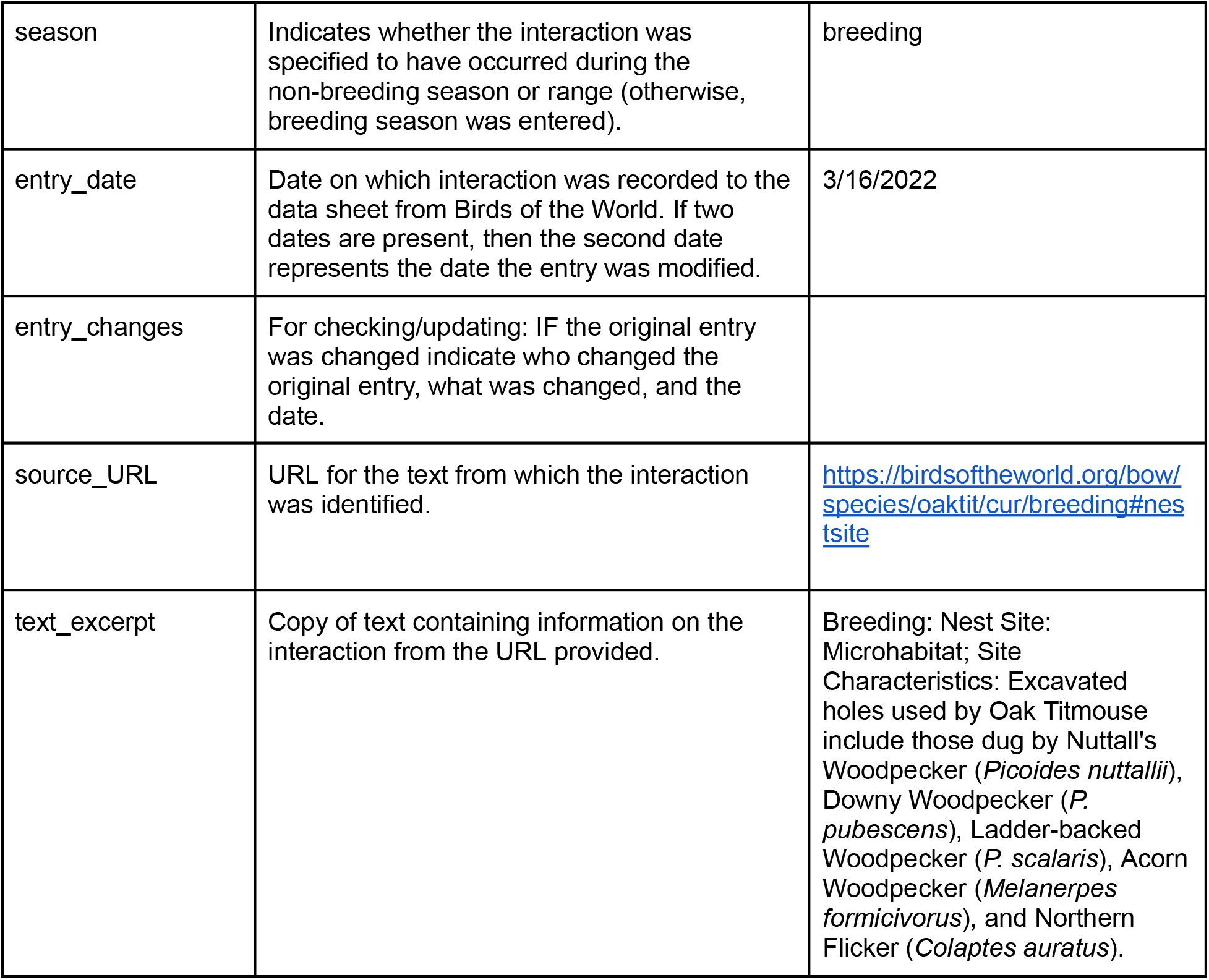
Data structure of the AvianMetaNetwork dataset. Included is one example text excerpt involving the focal taxa, Oak Titmouse (*Baeolophus inornatus*) and an interaction involving Nuttall’s Woodpecker (*Dryobates nuttallii*). References to the Clements/eBird checklist refer to the 2024 version ^47^.

| Column name | Description | Example |
| --- | --- | --- |
| taxa1_common | Common name of taxa1 according to Birds of the World (BOW) | Oak Titmouse |
| taxa1_scientific | Scientific name of taxa1 according to Clements/eBird checklist. If not able to be harmonized to Clements/eBird this name was left as recorded. | *Baeolophus inornatus* |
| taxa1_clements | Scientific name of taxa1 according to Clements/eBird checklist. (Note: In most cases this will be the same as taxa1_scientific unless the documented interaction provided taxonomic details not available in the Clements/eBird checklist.) | *Baeolophus inornatus* |
| taxa2_common | Common name of taxa2 according to Birds of the World (BOW) | Nuttall's Woodpecker |
| taxa2_scientific | Scientific name of taxa2 according to Clements/eBird checklist. If not able to be harmonized to Clements/eBird this name was left as recorded. (Note: In most cases this will be the same as taxa2_scientific unless the documented interaction provided taxonomic details not available in the Clements/eBird checklist.) | *Dryobates nuttallii* |
| taxa2_clements | Scientific name of taxa2 according to Clements/eBird checklist. | *Dryobates nuttallii* |
| interaction | Type of interaction occurring (see Table 1 for details). | commensalism |
| effect_tx1_on_tx2 | The effect of taxa1 on taxa2 as a result of the interaction (-1 = detrimental, 0 = neutral, 1 = beneficial) | 0 |
| effect_tx2_on_tx1 | The effect of taxa2 on taxa2 as a result of the interaction (-1 = detrimental, 0 = neutral, 1 = beneficial) | 1 |
| season | Indicates whether the interaction was specified to have occurred during the non-breeding season or range (otherwise, breeding season was entered). | breeding |
| entry_date | Date on which interaction was recorded to the data sheet from Birds of the World. If two dates are present, then the second date represents the date the entry was modified. | 3/16/2022 |
| entry_changes | For checking/updating: IF the original entry was changed indicate who changed the original entry, what was changed, and the date. |  |
| source_URL | URL for the text from which the interaction was identified. | https://birdsoftheworld.org/bow/species/oaktit/cur/breeding#nestsite |
| text_excerpt | Copy of text containing information on the interaction from the URL provided. | Breeding: Nest Site: Microhabitat; Site Characteristics: Excavated holes used by Oak Titmouse include those dug by Nuttall's Woodpecker (*Picoides nuttallii*), Downy Woodpecker (*P. pubescens*), Ladder-backed Woodpecker (*P. scalaris*), Acorn Woodpecker (*Melanerpes formicivorus*), and Northern Flicker (*Colaptes auratus*). |

For any interaction involving two bird taxa (a Taxon 1 and its interaction with a Taxon 2), recorders entered the interaction sub-type (Table 1), and a series of attributes relating to the interaction (Tables 1-2). The attributes of an interaction include: scientific and common names for each taxa, interaction sub-type, directional effects on each species, interaction timing (non-breeding or breeding), text excerpt and source, and notes. The interaction sub-type is a more specific form of eight interaction categories (competition, predation, commensalism, facilitation, parasitism, amensalism, mobbing, or other; Table 1). The effect (i.e., direction) of an interaction on each taxa is recorded as 0, 1, or -1 (neutral, negative or positive). Neutral relationships that do not fall strictly within the community ecology context of an interaction (sensu ^1,64–66^) are included for completeness; this includes interactions such as hybridization, co-occurrence, play, and other relationships that have unknown effects in terms of beneficial or detrimental impacts on taxa. When available, information on the timing of the interaction was noted in terms of the focal taxon’s non-breeding season or range; otherwise, breeding season was entered. If an interaction attribute was unclear from the BOW phrasing and a source was referenced, the original source was consulted. If the interaction was still unclear, a note was recorded in the “uncertain_interaction” column as an indicator to the data reviewer (see *Data Cleaning and Subsetting* below).

For each pairwise interaction, the text excerpt (including associated BOW page headings or figure caption) and accompanying BOW page URL were recorded. If it was difficult to interpret the interaction from the BOW text, the cited primary source was consulted, and noted. If an interaction between two specific avian taxa was described multiple times within a species account, or in a separate species account, each unique text excerpt received its own row. To ensure the data captured the most comprehensive set of interactions, all mentions of avian taxa in each Taxon 1 BOW account were considered, regardless of whether Taxon 1 was involved. In other words, if multiple species were involved in an interaction at one time (e.g., a group of species mobbing a potential predator), each pairwise interaction was recorded among taxa.

Throughout the project, we managed collaborative data entry and review using a combination of Google Sheets (for data entry and data edits) exported to CSVs for the L0 (raw) data, which were subsequently cleaned with R scripts to generate L1 (clean) data, that were further explored as L2 summaries (Figure 3). We stored interaction data in CSV format text files, grouped by species, using the git version control system ^67^ to track all changes and ensure data provenance.

### Data Cleaning and Subsetting

To create analysis-ready Level 1 data we implemented both manual and automated error checks to fix typos and ensure that all rows have the minimum necessary information (e.g., scientific names, interaction sub-type). Once a focal species’ interactions were input, the data sheet was checked and, if needed, corrected by a more experienced data recorder. All changes to the raw data sheet were indicated in an “entry_changes” column in the data sheet and tracked by GitHub.

Reviewed species data sheets were then concatenated into a master data sheet. Data were automatically checked for missing values in critical fields (e.g., scientific name, common name, interaction) and manually corrected where needed. To speed the processing of typos and other common errors (e.g., hyphens, capitalization), we generated a lookup table with common misspellings and their corrections and automatically processed these changes. We then harmonized taxonomy so that all common and scientific names followed the Clements/eBird 2024 checklist (see *Taxonomic Harmonization* below for details).

After harmonization, we used the regional species list to subset the data to only include focal taxa (i.e., “Taxa 1”) from within the continental United States, Alaska, and Canada (while retaining records of interactions of focal taxa with non-focal taxa).

### Taxonomic Harmonization

With increasing knowledge of species’ evolutionary histories and movements, bird taxonomy has experienced many changes in recent years. Many species that were once considered distinct have now been “lumped” together, while other species have been “split” into multiple groups. Still more species groups have been moved into entirely new genera. These changes can create challenges when synthesizing past literature. For instance, a species that was “split” may have fewer documented interactions, not because the individual birds are not interacting *per se*, but because all historical observations prior to the split refer to the species by their previous, aggregated name. All together, these changes necessitate careful consideration of name change history and choice of taxonomic authority.

Throughout the workflow, all taxa names were aligned with the Clements/eBird 2024 checklist for taxonomic harmonization ^47^, and any discrepancies were changed in code (Figure 3 “Clean database”). To harmonize taxonomy for scientific names at the species and subspecies levels in the data that did not exactly match names on the Clements/eBird 2024 checklist (401 of 1,989 taxa), we ran semi-automated checks to discover the correct alignment, followed by a transparent, manual name exchange. This process discovered both those taxa with updates but also those with typographic errors from the data entry process. First, we used fuzzy text search on scientific names that returned a probable match from our current checklist and the Jaro-Winkler distance ^68^ from the stringdist package to discover corrections for typographical errors ^69^. We manually checked the results to determine our high-confidence threshold distance score of 0.95/1.0 that always included correct matches (132 of 401 taxa). For the remaining 269 unresolved names, the scientific names either had significant typos or more likely had a taxonomic change. However, frequently the common names remain the same, so we re-applied our fuzzy text search to match common names in the database with our current checklist. After manually determining a high-confidence threshold of 0.94/1.0, we matched 225 scientific names, leaving 50 remaining. We used taxonomic entries from GBIF via the taxaDB R package ^70^ to identify known synonyms. However, GBIF avian taxonomy is not up to date with Clements/eBird 2024, resulting in limited capacity for taxaDB to match as many taxa (11 taxa). Finally, for the remaining discrepancies (including genus-level or higher-level taxa) we manually evaluated the scientific names, referring to the Clements/eBird 2024 checklist ^47^, Avibase ^60–62^, the 2024 BBS species list ^63^, and BOW ^57^ to create a list of manual taxonomic corrections. These discrepancies could be full scientific names we could not match (for example newly split *Bulbucus ibis* to *Ardea ibis/coromanda*), hybrids (for example, *Anser indicus x Branta leucopsis*), genus-level taxa (e.g., *Accipiter sp*.), or taxonomic changes (*Chen rossii* to *Anser rossii*).

### Data Record

AvianMetaNetwork consists of two primary files, archived in CSV format with the Environmental Data Initiative. The first file (AvianMetaNet_InteractionData_EDI.csv) consists of all documented instances of bird-bird interactions. Each row in the csv is a unique instance of a documented interaction (Table 2). Column names include the scientific and common names of each of the two taxa, the equivalent Clements name for each of the two taxa, the interaction type (Table 1), the effect of taxa 1 on taxa 2 as a result of the interaction, the effect of taxa 2 on taxa 1 as a result of the interaction, the season, a URL and a text excerpt from Birds of the World containing the evidence of the interaction, the date of the entry, and a description of any manual changes made to the entry after initial coding.

The second file (spp_clem_in_amn_cac.csv) is the complete taxonomic checklist of all taxa in AvianMetaNetwork. This file is derived from the Clements/eBird 2024 checklist described above and contains all of the original columns in the checklist, but is subsetted to only include the taxa in AvianMetaNetwork. We have added an additional binary column “canada_ak_conus” to indicate whether we considered a taxa to be a Canada, Alaska, or Coterminous United States taxa (TRUE) or not (FALSE).

### Data Overview

We provide 25,907 total records documenting pairwise interactions. As the data contain multiple records of the same pairwise interaction (i.e., multiple rows may contain multiple accounts of the same taxa pair interacting in the same way, but with different text sources), the unique number of pairwise interactions is smaller, totaling 18,431. The most common interaction type is predation (6,504 total and 4,786 unique records), whereas the least common interaction type is amensalism (73 total and 71 unique interactions; Figure 1).

The majority of unique interactions (76%) occur between focal taxa. In the process of entering bird-bird interactions for a focal species, data recorders often encountered an interacting species that was not on the regional species checklist. In addition to the 731 focal taxa, 1,258 more taxa were found to interact with these species outside of the study region (i.e., while a focal species was in migration or in a part of the focal species’ range that was outside the study region). Aside from records involving a few vagrants/accidental species with ranges fully outside the Western Hemisphere (e.g., Carrion Crow (*Corvus corone*), White-tailed Eagle (*Haliaeetus albicilla*), and Whinchat (*Saxicola rubetra*)), most of these 1,258 taxa are native to Central America. For instance, the Ruby-throated Hummingbird (*Archilochus colubris*, a migratory species that breeds in the eastern United States) competes with non-migratory hummingbird species (e.g., the Sparkling-tailed Hummingbird, *Tilmatura dupontii*) while in their non-breeding range, which spans southern Mexico and northern Central America. Similarly, the Cactus Wren (*Campylorhynchus brunneicapillus*, a non-migratory species that resides in the southwestern United States and Mexico) communally roosts with other congeneric wrens, namely, Boucard’s Wren and Spotted Wren (*C. jocosus* and *C. gularis*, respectively) where their ranges overlap in central and southern Mexico, outside the continental United States.

We provide three examples of the networks built upon three focal species: Oak Titmouse (*Baeolophus inornatus*), Nuttall’s Woodpecker (*Dryobates nuttallii*), and Northern Pygmy-Owl (*Glaucidium gnoma*). These three species co-occur year-round in the dry, open, oak-pine and oak woodlands along California’s Coast Ranges and the western foothills of the Sierra Nevada Mountains ^71–73^. Within these habitats, the three focal species exhibit several interactions with each other. Nuttall’s Woodpeckers excavate cavities for nesting, which are often used by secondary cavity nesting bird species in subsequent years, including the Oak Titmouse. These species also compete for nesting habitat, with Nuttall’s Woodpeckers usually dominating Oak Titmouse pairs that nest within the same tree. Northern Pygmy-Owls are small, predatory birds that take a variety of prey, including Nuttall’s Woodpeckers. Oak Titmouse are also susceptible to predation from the Northern Pygmy-Owl, and respond through mobbing, often in association with other small birds. We constructed each species network by filtering for interactions among each focal species and the species they directly interact with, then using the graph_from_data_frame function in the R package igraph (version 2.2.1, ^74–76^). Each network was visualized using the ggraph R package (version 2.2.2, ^77^).

### Technical Validation

To extract species interaction information from Birds of the World (BOW) for each focal species, data recorders were trained by senior personnel on the project. As part of the data entry training, all recorders worked on the same BOW species account to calibrate their data entry. Data recorders and senior personnel then met to discuss any discrepancies among recorders and against the species’ final validated data (entered data that was checked by a more experienced data recorder). To ensure the quality and interoperability of the AvianMetaNetwork data, we used both manual and automated validation procedures. All data sheets were entered initially by a data recorder and then manually checked by a more experienced data recorder. Manual checking involved re-referencing BOW accounts to verify that all appropriate fields were entered correctly (see *Data Entry* for further details).

Uncertainties in interactions and other interpretations of the text were resolved through discussion among senior personnel and changes to the data sheets were tracked using Google Docs and GitHub. The checking of initial data entries by a more experienced recorder, and regular project-wide meetings to discuss interpretations, ensured that any discrepancies were considered by at least two individuals. Through this process, experienced recorders discovered errors or omissions in 13.5% of the rows, and corrections were applied, with the reason noted in the “entry_changes” column. Throughout the process of data entry and checking, independent sources outside BOW were used to verify taxa names (Avibase v8.17, Clements/eBird 2024 checklist) as well as the BOW species’ accounts of the interacting species and original sources cited within a BOW account.

Once completed and manually validated, individual species interaction data sheets were further cleaned via automated checks to standardize data classes (e.g., numeric, string) and remove readily identifiable errors (e.g., trailing spaces in strings) prior to data aggregation into a single dataframe. Finally, both automated and manual steps were taken to correct typos and harmonize taxonomy in the data (see *Taxonomic Harmonization* for details).

### Usage Notes

We present these data at the most comprehensive level to enable users to subset the network to the desired level of taxonomic resolution. Users can further subset the data to a given taxonomic resolution by aligning with the subsetted Clements/eBird checklist that we provide alongside the main dataset in the EDI repository. It should be noted that when interactions are reported at a coarser taxonomic resolution (e.g., family-level), it is not possible to determine whether the provided information indicates that a given focal taxa was documented to be interacting with a single species within a family or with all members of a family. For example, if a Peregrine falcon (*Falco peregrinus*) has a predation interaction with “Columbidae”, it is not possible to resolve whether the Peregrine has been documented to predate upon all members of the family or just a single or a few species within this family.

One limitation of the approach is the reliance on BOW accounts for interaction information. Although BOW is the most comprehensive collection of avian natural history ^57^, some BOW species accounts contain limited information (due to lack of knowledge, out of date accounts, or a combination thereof). As such, some sets of interactions may be incomplete. To fill in gaps and ensure improved completeness, a future direction of this project will expand the search beyond BOW and perform a more comprehensive search via API access to literature databases (PubMed Central, Google Scholar, and Web of Science), using species’ current and past common and scientific names (defined by the current Clements/eBird checklist ^47^ and Avibase ^78^).

The data presented here represent a snapshot of an ongoing research effort to systematically create a network of all known bird interactions in the world. In the interest of open science, we have made our data cleaning and analysis workflow for the project publicly available at the AvianMetaNetwork public project GitHub site ^79^; the specific data and code used in this paper covering the region of the continental United States and Canada is included in an EDI repository (see *Data and Code Availability*). The repository includes R and Quarto Markdown (QMD) files, named in 8 successive steps, which should be run in order (steps 1 and 2 are within /R/L0, steps 3-6 are within /R/L1, and steps 7 and 8 are within /R/L2). The final two QMD files (7_figure_processing and 8_summary_vignette) also include their html output, and are helpful for visualizing the data and reproducing the figures in this paper. We note that script 7_figure_processing_vignette.qmd is a walkthrough of the data cleaning and formatting needed to produce the figures in this paper from the AvianMetaNetwork data, and also contains the function definitions that are used in plotting. The script 8_summary_vignette.qmd contains summary statistics and plotted figures.

## Data Availability

Data are archived with the Environmental Data Initiative (EDI). (See below for access information. Data are currently embargoed until data paper publication at which time a DOI will be assigned). Publicly available data used in the creation of AvianMetaNetwork include: the Clements/eBird 2024 checklist ^47^, Avibase v8.17 regional checklists for “US48”, “Canada” and “Alaska” ^60–62^, and the North American Breeding Bird Survey species list ^63^. As the AvianMetaNetwork project is currently expanding beyond the region covered in this paper, we direct readers to our public GitHub repository ^79^ to view the latest progress. To prevent accidental misuse and/or misinterpretation of raw data while still adhering to the principles of open science, we store all in-progress interaction data in a separate, private GitHub repository while keeping code, metadata, and species lists publicly accessible in the aforementioned GitHub repository. EDI embargoed data and code repository: Zarnetske, P.L., P. Bills, K. Kapsar, L. Mansfield, and E. Parker. 2026. The AvianMetaNetwork: biotic interactions among birds of the continental United States and Canada ver 2. Environmental Data Initiative. https://doi.org/10.6073/pasta/9bc99f67618359b2d9a6770eff22664a.

## Code Availability

Code is archived with the Environmental Data Initiative (EDI). (See *Data Availability* for access information).

## Acknowledgements

We thank Michigan State University (MSU) undergraduates who have been involved in compiling additional data on this project: Arpita Nayak, Minali Bhatt, Erik Ralston, Jordan Zapata, Elaine Hammond, Ava Fountain, Ann Joseph, Maddie Andreatta, Sarah Pecis, Olive Graves, Vivian Smith, Liz Bauer, Addison Hoddinott, Elliot Palmer, and Jaime Soehl. We also thank the Cornell Lab of Ornithology for providing the Birds of the World Online resource and for helpful feedback on this project (including Brian Sullivan, Pam Rasmussen, Jeff Gerbracht, Eliot Miller, Laura Kammermeier, Paul Rodewald). We also thank Vincent Miele, Stéphane Dray, David Skelly, Mark Urban, Jenna Baljunas, and Minyoung Lee for their helpful insights.

